# *takeout* gene expression is associated with temporal kin recognition

**DOI:** 10.1101/2023.06.12.544599

**Authors:** Ahva L. Potticary, Elizabeth C. McKinney, Patricia J. Moore, Allen J. Moore

## Abstract

A key component of parental care is avoiding killing and eating your own offspring. Many organisms commit infanticide but switch to parental care at the time when their own offspring would be expected, known as temporal kin recognition. It is unclear how such indirect kin recognition is so common across taxa. One possibility is that temporal kin recognition can be achieved by altering a simple mechanism, such as co-opting the regulation of timing and feeding in other contexts. Here we determine whether *takeout*, a gene implicated in coordinating feeding, influences temporal kin recognition in the roundneck sexton beetle, *Nicrophorus orbicollis*. We find that *takeout* expression is not associated with non-parental feeding changes resulting from hunger, or in the general switch to the full parental care repertoire. However, beetles that accepted and provided care to their offspring had a higher *takeout* expression than beetles that committed infanticide. Together, these data support the idea that the evolution of temporal kin recognition may be enabled by co-option of mechanisms that integrate feeding behaviour in other contexts.

## INTRODUCTION

Parenting is expensive, and thus a variety of mechanisms to avoid caring for unrelated offspring have evolved. Many organisms commit infanticide of conspecific young but switch to parental care at the time when their own young would be expected, known as temporal kin recognition (Beecher et al., 1981; Elwood, 1994; Müller & Eggert, 1990; Perrigo et al., 1992). The time lag between the cue that “starts the clock” and the behavioural shift to parental care can be quite long. For example, male mice are infanticidal until 18-20 days after ejaculation, after which they show parental care to young (Perrigo et al., 1992). Temporal kin recognition often works in the absence of conspecific phenotypic cues; indeed, parents presented with young at the correct time and place will even accept and feed young of other species (e.g., Benowitz et al., 2015; Davies, 2000; Elwood, 1994). Temporal kin recognition is a short-term biological timing problem - that is, it can occur over several days, is reversible and not seasonal - and has been linked to circadian rhythms across taxa (Oldekop et al., 2007; Perrigo et al., 1992). For this reason, it has been suggested that co-option of the mechanisms that regulate feeding and timing in other contexts are co-opted to enable the evolution of temporal-based kin recognition (Moore et al., 2010).

Here, we investigate whether a *takeout*, a gene known to be associated with coordinating feeding behaviour through an association with circadian rhythms (Meunier et al., 2007; Sarov-Blat et al., 2000), has been co-opted to facilitate temporal kin recognition in the burying beetle *Nicrophorus orbicollis*. Two RNA-seq studies of different species of burying beetle comparing gene expression in a parenting state with a non-parenting state identify *takeout* as one of the genes differentially expressed (Moss et al., 2022; Parker et al., 2015). Thus, *takeout* appears to play some role in burying beetle parenting although the transcriptomic studies could not identify the specific function given the broad comparison. In insects, *takeout* gene expression is involved in pathways which convey temporal and nutritional information to influence feeding activity (Sarov-Blat et al., 2000), has been linked to attraction or aversion to chemosensory cues (Guo et al., 2011), and is often located in areas that correspond to chemosensory functions such as the antennae (Fujikawa et al., 2006). *takeout* mRNA can cycle in the absence of light in *Drosophila* (So et al., 2000), although *takeout* can also be influenced by light and dark cues (Benito et al., 2010). We hypothesized that because parenting requires a switch from eating offspring to feeding offspring underground, and because this switch occurs at a very specific time in burying beetles, this switch will involve *takeout*.

Burying beetles require small animal carcasses to breed, and once a suitable carcass is located, the female lays eggs in the soil nearby and the parents bury and prepare the carcass as a food resource for their offspring (Scott, 1998). Upon hatching, larvae crawl to the carcass where they are fed regurgitated carrion by the parents. Burying beetles identify their larvae temporally; parents of both sexes will commit infanticide of any larvae that arrive too early, and will only accept and feed larvae that arrive at the time where their own offspring can be expected (Müller & Eggert, 1990; Oldekop et al., 2007). However, parents will accept any larvae that arrive at the correct time, even larvae of other burying beetle species (Benowitz et al., 2016). Temporal kin recognition in *Nicrophorus* can be manipulated by changing light schedules (Oldekop et al., 2007); thus, the mechanism that influences the timing of temporal kin recognition is still sensitive to photic inputs despite *Nicrophorus* breeding underground, which hints at co-option of mechanisms involved in daily cycling. Intriguingly, the embryonic periods or time to larval acceptance for many *Nicrophorus* occur in increments of approximately 24-hours; for example, larvae are accepted at approximately 48 h post egg-laying in *Nicrophorus mexicanus* and *N. vespilloides* (Anduaga, 2009; Oldekop et al., 2007), 72 h in *N. quadripunctatus* (Takata et al., 2015), and 96 h in *N. orbicollis* (this study). Some burying beetles only bury the carcass under the leaf litter, like *N. vespilloides*, while others such as *N. orbicollis* bury the carcass multiple centimeters below the soil surface (Scott, 1998; Wilson & Fudge, 1984). For these reasons, mechanisms involved in both feeding and timing, that can cycle in the absence of light cues, may be particularly likely to enable the evolution of temporal kin recognition in *Nicrophorus* (Moore et al., 2010).

Our hypothesis that *takeout* has been co-opted to influence temporal kin recognition makes several predictions. First, *takeout* should be associated with non-parental contexts that have been previously linked to *takeout* in other species. Second, *takeout* should not be associated with parental care in general, such as preparation of the carcass. Third, *takeout* expression should differ between parents that accept larvae and those that commit infanticide independently of any role in non-parental contexts or in general parenting. To test these predictions, we first experimentally induce hunger in beetles to determine the effect of food deprivation on *takeout* expression. We next measure *takeout* expression during the development of parenting behaviour. We then collect parents both as they show natural variation in temporal kin recognition and experimentally manipulate when parents expect their larvae to arrive to determine whether *takeout* expression differs between parents that accept or commit infanticide. We find that *takeout* expression is associated with the likelihood that parents will accept their larvae but find little evidence for an association between *takeout* and non-parental roles or the development of parenting.

## MATERIALS AND METHODS

### Ethical Note

We follow the ASAB/ABS guidelines for the treatment of animals in behavioural research and describe the origin and housing conditions for all beetles used in this study below. We designed housing with the goal of reducing stress and maximizing the animals’ welfare. We complied with the regulations in the USA for experiments on invertebrates.

### Insect Colony and Husbandry

*Nicrophorus orbicollis* used in this study were part of an outbred colony that we supplemented regularly with beetles from Whitehall Forest in Athens, GA (Potticary et al., 2023). We maintained all virgin beetles in an incubator (Percival, model no. I41VLC8, Perry, IA) at 22 ± 1 °C on a 14:10 hour light:dark cycle to simulate summer breeding conditions. All individuals used for this study were bred in the laboratory to ensure that parentage, age, and rearing conditions were known and standardized. We housed beetles individually at larval dispersal in 9-cm-diameter by 4-cm-deep circular compostable deli containers (EP-RDP5, Eco-Products) filled with ∼2.5 cm of potting soil (Performance Organics All Purpose In-Ground Soil, Miracle-Gro). Once we placed larvae into individual containers, they had no further social interactions with other burying beetles until we allocated them into the behavioural or non-parental treatments. We moved beetles that were allocated to the parental treatments (development of parenting and temporal kin recognition) to an incubator under constant darkness to simulate underground breeding, while beetles in the non-parental treatment (hunger vs fed) were kept in the same incubator as other virgins. We defined individuals as age Day 1 on the day of eclosion. We fed beetles *ad libitum* organic ground beef twice a week following ecolsion. For all treatments, we randomly mated focal virgins aged Day 13 - 19 post-eclosion with a non-relative.

### Gene Identification

We used Geneious Prime (v. 2022.1.1) to blast the *Nicrophorus orbicollis* transcriptome (Won et al., 2018) for putative takeout homologues of *D. melanogaster* (Q9VBV3) and *N. vespilloides* (XP_017773976). To visualize protein conservation across *takeout* copies, we aligned protein sequences from *Drosophila melanogaster* (Sarov-Blat et al., 2000), *Nicrophorus vespilloides, Locusta migratoria* (Guo et al., 2011), *Reticulitermes speratus* (Fujiwara et al., 2023), *Bombyx mori* and *Anopheles gambiae* (Saito et al., 2006), and using Clustal Omega (https://www.ebi.ac.uk/Tools/msa/clustalo/) and produced plots with Javalview (Waterhouse et al., 2009).

### Takeout Expression in Non-parental Contexts

To test the hypothesis that *takeout* is associated with hunger in non-parental contexts, we experimentally manipulated food availability for female and male beetles. We selected virgin beetles Day 13 – 19 post-eclosion and haphazardly allocated them to a hunger treatment (no feeding for one week), and a fed treatment (*ad libitum* feeding for one week). The goal of this manipulation was to induce hunger without resulting in starvation or mortality. We selected a week for the length of the manipulation because *Nicrophorus* beetles can survive for multiple weeks without food (Steiger et al., 2007), and body mass loss can occur in *N. orbicollis* over short periods of time from similar periods of withholding food (Trumbo & Robinson, 2004; Trumbo & Xhihani, 2015). We conducted this experiment with n = 14 females and males in each treatment, although three female beetles died during the experiment (n = 2 “hunger” females and n = 1 “fed” female).

### Takeout Expression and Parenting

#### *Takeout* and the Development of Parenting

To test the hypothesis that *takeout* is associated with the development of parenting, defined as the days elapsed since egg-laying and specifying the period where parents transition from a non-parenting state to a state when larvae may be accepted, we collected female and male beetles at 0 (eggs laid), and then at 24-, 48-, 72-, and at 96-hours post-egg laying, when larvae should first appear at the carcass. We mated and allocated pairs to breeding boxes with approximately 2 cm of potting soil. We checked breeding boxes every hour from 8 am – 5 pm for eggs; we only included set-ups where the breeding box was observed with no eggs one hour and with at least one fertile egg the next hour. After eggs were laid, we allocated individuals to treatments (0, 24, 48, 72, and 96) and collected beetles relative to the time that their eggs were first observed (e.g., if eggs were first observed at 9 am, adults in the 24-hour treatment were collected at 9 am the subsequent day). We collected n = 10 of both sexes for each treatment.

#### *Takeout* and Temporal Kin Recognition

##### Manipulation of larval arrival time

To test the hypothesis that *takeout* specifically influences the transition from infanticide to acceptance and feeding of larvae, we performed an experiment where we manipulated the temporal cues used by parents to anticipate offspring arrival to sample adults as they committed infanticide of the larvae or accepted and fed offspring. After we paired beetles with a mouse and observed eggs, we moved the focal female and the brood ball to a new breeding box and placed a small piece of beef in the original breeding box with the eggs to attract the larvae and checked for hatch 1-3 times a day. We kept the focal male separate from the female and brood ball for 24 hours and then returned to the male to the breeding box with the female. Parents were separated and reunited to “restart the clock” on temporal kin recognition, such that the pairs’ own larvae from the first batch would arrive before parents expected them.

After reuniting the male and female we again checked breeding boxes daily for eggs. For both treatments, we provided the breeding pair with ten larvae, at least 5 of which were their offspring from the first clutch of eggs and were supplemented with same-age larvae from other clutches if insufficient numbers were available from the focal pair’s clutch, as *N. orbicollis* cannot identify their own larvae (Benowitz et al., 2016; Oldekop et al., 2007). Following the addition of larvae, we observed the focal individuals for at least thirty minutes to detect acceptance or rejection of larvae following established protocols (Potticary et al., in press). If the focal individuals did not respond in this time, pairs were checked every 30 minutes to identify the onset of acceptance or infanticide. In some cases, one parent would accept larvae while the other parent committed infanticide; in these cases, the male and female were sampled and categorized separately according to the phenotype that they exhibited. For the Infanticide treatment, focal individuals were collected while the focal individual was attacking/eating larvae. As the larvae were provided prior to when parents expected to find them, it often took extended periods of time for parents to detect and respond to larvae. If the parents did not eat the larvae within an hour of larval placement, the carcass was checked every 15-30 minutes until aggression towards larvae was observed. For the Accept treatment, focal individuals were collected immediately after the parent made antennal contact with the larvae and a feeding event was observed. This manipulation often led to a discrepancy in the temporal cues used by males versus females, as females base their window of larval acceptance on when eggs are laid (Oldekop et al., 2007), and males likely base their window on timing and frequency of female acceptance of mating (Potticary, pers. obs.). Thus, to reach an adequate sample size, we removed the female after the first batch of eggs were laid, which increased the likelihood that males would accept larvae. We collected n = 20 for both sexes in both treatments.

##### Natural variation in temporal kin recognition

We took advantage of natural variation in temporal kin recognition to test the hypothesis that variation in acceptance or infanticide of larvae is associated with variation in *takeout*. To do this, we followed the methods above but altered the procedure such that the focal beetles were provided larvae at the correct time. Specifically, after we detected eggs, we allowed the pair to remain in the breeding box for 24-36 hours to ensure that egg-laying was completed. Then, the focal individual was moved with the brood ball to a new breeding box to ensure that the larvae did not arrive at the carcass naturally, and the non-focal parent was removed. Once we detected larvae and/or 96 hours had elapsed since we detected the first fertile egg in the original breeding box, we provided focal parent with a mixed brood of five larvae to determine whether parents accepted or committed infanticide using the same behavioural methods described above. However, this manipulation also resulted in abandonment, where the parent ceased attending to the carcass, which resulted in significant decay and fungal growth on the carcass, and/or the parent failing to feed larvae despite contact. As both infanticide and abandonment both constitute a lack of parental care towards larvae, we categorized infanticide and abandonment as rejection. After observation of acceptance or rejection of larvae (including both infanticide and abandonment), we collected the heads of focal individuals to determine how natural variation in temporal kin recognition was associated with *takeout* gene expression. We collected n = 19 females and n = 25 males for this experiment.

### Gene Expression Analysis

We collected samples by removing the beetles’ heads with dissecting scissors and then immediately flash froze them in liquid nitrogen (Roy-Zokan et al., 2015). We then stored samples at -80 °C until RNA extraction. Samples were homogenized in liquid nitrogen using a mortar and pestle (cat. no. 60310, CoorsTek, Golden, CO, USA). Following homogenization, we extracted RNA from each sample using Qiagen’s RNAeasy Lipid Tissue Mini-Kit (cat. no. 74106) following (Roy-Zokan et al., 2015). We also treated samples with DNase I (Qiagen) on the column according to manufacturer’s instructions to help ensure minimal genomic DNA contamination. We quantified RNA using a Qubit 2.0 Fluorometer (Invitrogen Corporation, Carlsbad, CA, USA) using the RNA Broad Range protocols per manufacturer’s instructions. We synthesized cDNA with Quanta Biosciences qScript reverse transcriptase master mix (Quanta Biosciences, Gaithersburg, MD, USA) following the manufacturer’s instructions from 500 ng total RNA. RNA was stored at -80 °C and cDNA was stored at -20 °C.

We performed quantitative real-time PCR (qRT-PCR) primers from the PCR-validated consensus sequences for our gene of interest and an endogenous control gene (*gapdh*) following methods of Cunningham et al. (2014) and using transcriptome of Benowitz et al. (2017). We performed qRT-PCR with Roche LightCycler 480 SYBR I Green Master Mix using a Roche LightCycler 480 (Roche Applied Science, Indianapolis, IN, USA) following the manufacture’s protocol with triple technical replicates of 10 μl reactions. Primers were at a working concentration of at 1.33 μmol/l. Annealing temperature was 60 °C during the amplification cycles. We established the stability of the endogenous reference gene by determining that *gapdh* did not vary across the sexes or experimental treatments. Additional information suggested by the Minimum Information for Publication of Quantitative Real-Time PCR Experiments (MIQE) guidelines can be found in Appendix 1.

### Statistical Analyses

All statistical analyses were performed using JMP PRO (v. 16.0.0, http://jmp.com) and all figures were produced in SigmaPlot (v. 14.5, http://www.sigmaplot.co.uk). Results are means ± SE. We used the ΔΔ*C*_T_ method (Livak & Schmittgen, 2001) to convert raw expression data to normalized relative expression values. For all analyses, we used the treatment group that occurred earliest as the comparison group. Females and males were analyzed together for all experiments, as previous research has found sex-biased expression of *takeout* (Dauwalder et al., 2002; Vanaphan et al., 2012) and thus there is an *a priori* expectation for an interaction between *takeout* and sex. All statistical tests conducted in this study were two-tailed, as there was no *a priori* expectation for the direction of associations between variables. We analyzed the manipulation of fed vs. hungry beetles, the development of parenting, and the experimental manipulation of Infanticide vs. Acceptance using two-way ANOVA with sex and treatment as factors. We analyzed *takeout* expression in the experiment of natural variation in infanticide versus reject using t-tests; for this experiment, only male data were included as only two females rejected larvae when larvae were presented at the correct time.

## RESULTS

### Gene Identification

Plots illustrating sequence homology between *N. orbicollis, N. vespilloides, L. migratoria, R. speratus, D. melanogaster*, and *A. gambiae* indicate moderate conservation between the different species (**Figure 1**). The *N. orbicollis takeout* studied here has a percent identity of 25.22% with the *D. melanogaster takeout*. This is comparable to the reported percent identities for *takeout* genes with the *D. melanogaster takeout* across species, including *N. vespilloides* (25.74%), *L. migratoria* (24.47%), *R. speratus* (RsTO1: 25.74%, RsT02, 24.69%), and *A. gambiae* (AgTOL1: 24.08%, AgT02: 40.0%).

**Figure 1.**
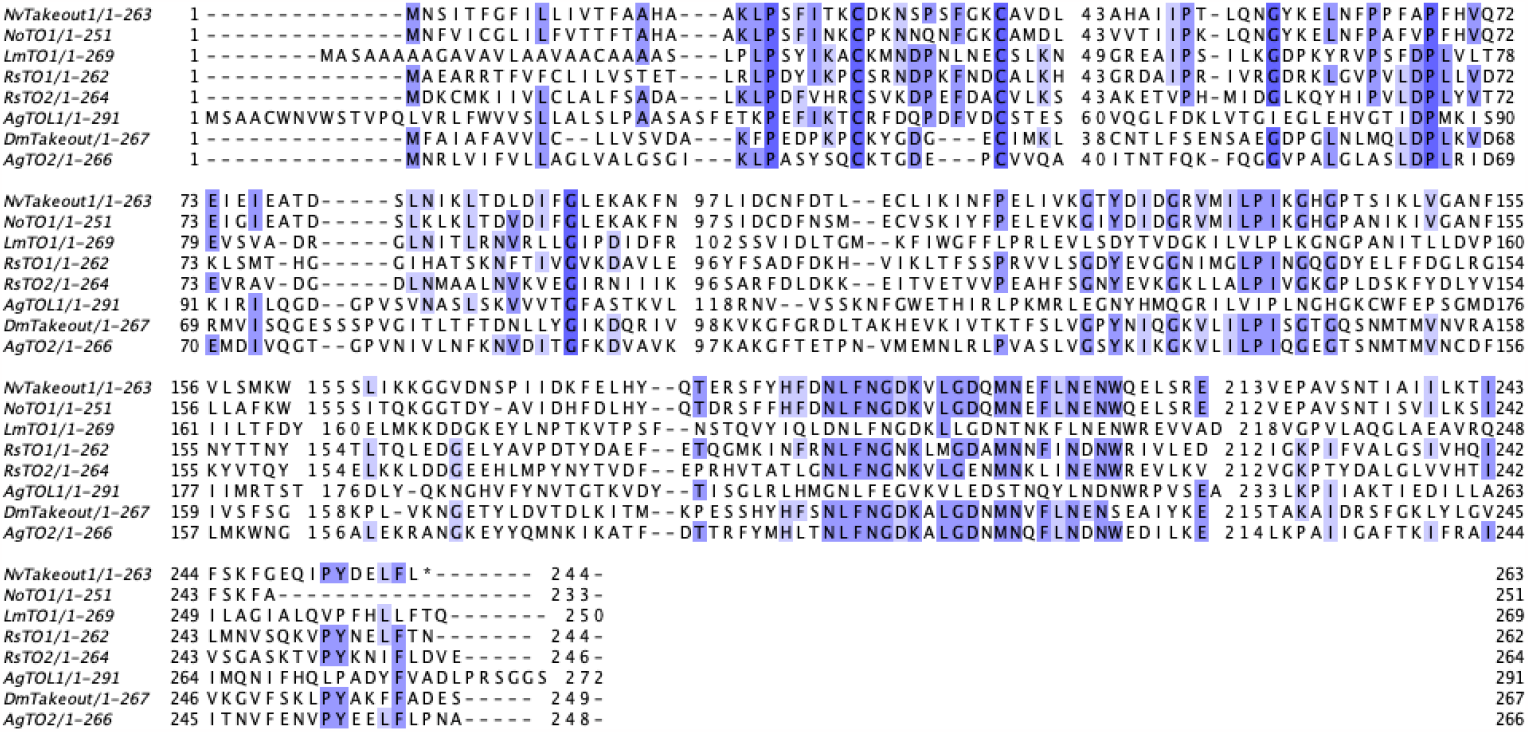
Amino acid alignment of *takeout* family. Amino acid alignment of members of *takeout* family in *N. vespilloides, N. orbicollis, L. migratoria, R. speratus, D. melanogaster*, and *A. gambiae*. Shaded regions represent > 40% similarity among sequences.

### Takeout Expression in Non-parental Contexts

There was no statistically significant difference in *takeout* expression between beetles that were fed versus hungry (F_1,50_ = 0.28, P = 0.60), the sexes (F_1,50_ = 1.74, P = 0.19), nor was there an interaction between treatment and sex (F_1,50_ = 1.40, P = 0.24; **Figure 2a**).

**Figure 2.**
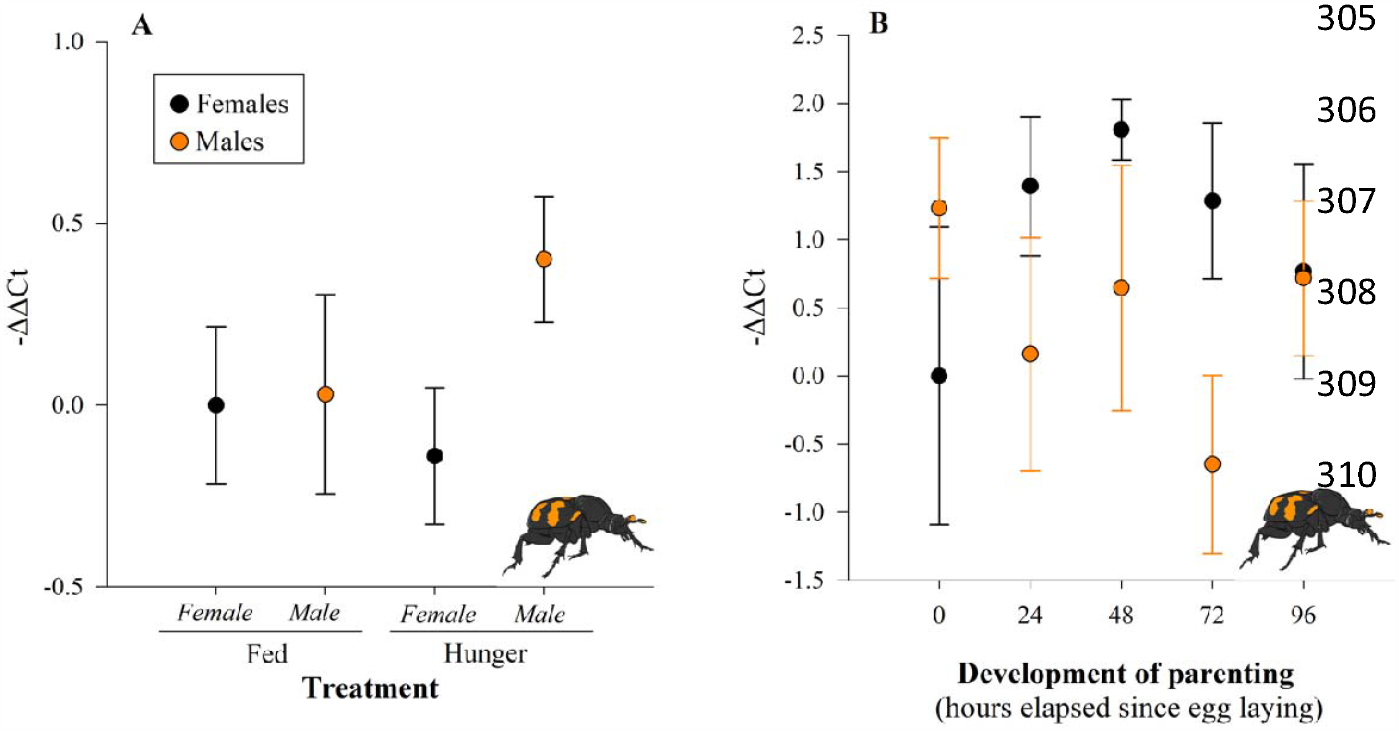
*takeout* is not associated with non-parental contexts or development of parenting. Closed circles represent females, while orange circles represent males. There was no association between whether beetles were (***A***) well-fed or hungry (females: n= 13 fed, n = 12 hungry; males: n = 14), nor (***B***) with the development of parenting in either female or male beetles (n = 10 for each treatment). Results are mean ± SEM.

### Takeout Expression and Parenting

We did not find an association between *takeout* and the development of parenting. *Takeout* expression was not associated with the development of parenting (F_4,94_ = 0.43 P = 0.79), sex of the beetle (F_1,94_ = 1.1.52, P = 0.22), there was no interaction between development of parenting and sex (F_4,94_ = 2.12, P = 0.20; **Figure 2b**).

*Takeout* was associated with temporal kin recognition in both females and males. In the experimental manipulation of larval arrival time, parents that accepted their larvae had higher *takeout* gene expression than beetles that committed infanticide of their own larvae (F_1,76_ = 5.03, P = 0.03; **Figure 3a**), and females generally showed higher *takeout* expression than males (F_1,76_ = 6.04, P = 0.02), but there was not a statistically significant interaction between treatment and sex of the caregiver on *takeout* expression (F_1,76_ = 0.25, P = 0.62). We found a similar pattern in the natural variation of accept versus reject; males that accepted had higher *takeout* expression than males that rejected their larvae (t_1,23_ = -2.12, P = 0.05; **Figure 3b**). There was an insufficient number of female beetles that naturally rejected their larvae to analyze. Male beetles showed more variation in accept versus reject when they were presented with larvae at the correct time (males: n = 15 accept versus n = 10 reject) than females (n = 17 accept versus n = 2 reject).

**Figure 3.**
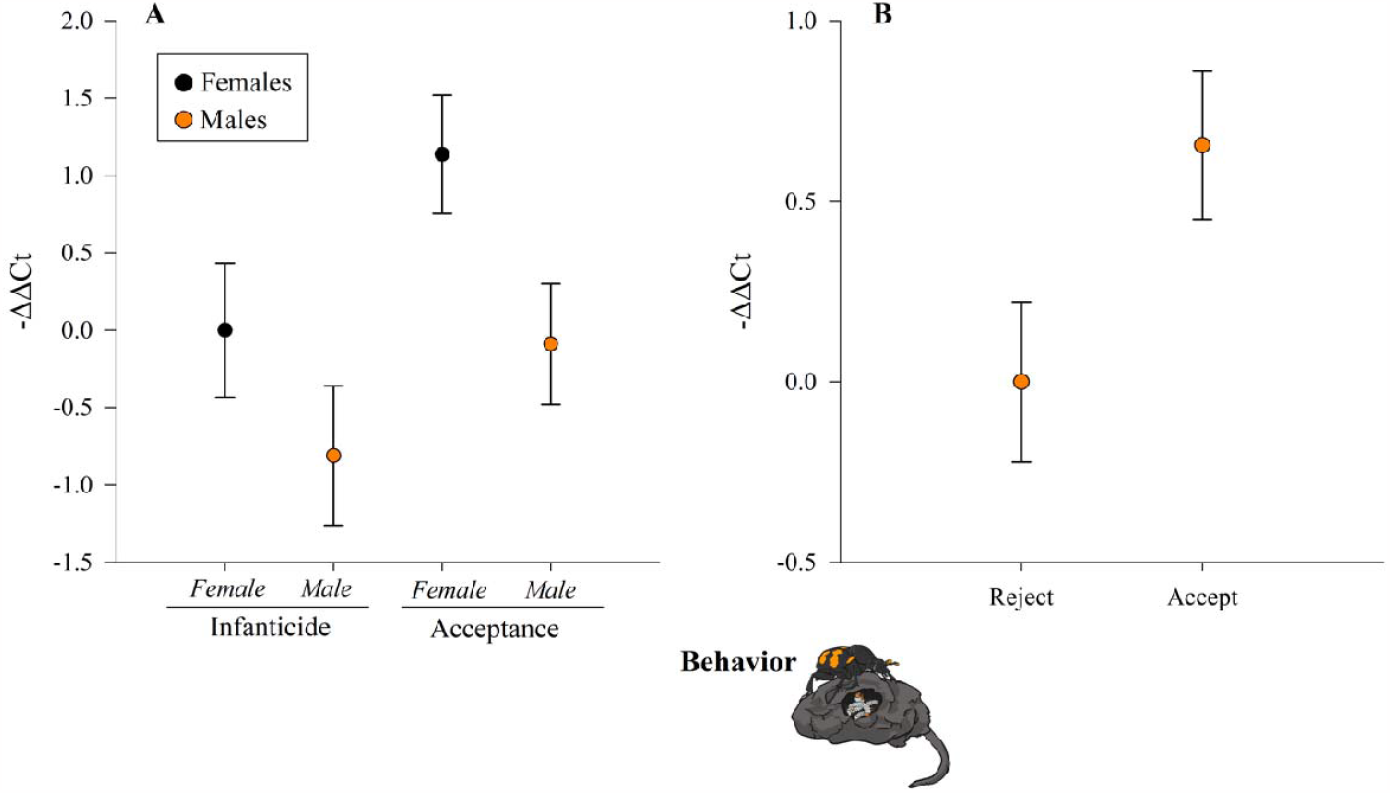
*takeout* is associated with temporal kin recognition. Closed circles represent females, while orange circles represent males. (***A***) Female and male parents that accepted their larvae had higher takeout than beetles that committed infanticide, and females generally showed higher takeout expression than males (n = 20 for each treatment). (***B***) Males showed natural variation in infanticide when presented with larvae at the correct time, and for these males, those that accepted their larvae also showed higher takeout expression than those that rejected their larvae (n = 10 reject, n = 15 accept). Results are mean ± SEM.

## DISCUSSION

Evolution of complex behaviours like temporal kin recognition can involve the co-option of behavioural precursors (Tallamy, 1984; West-Eberhard, 2003) and thus the mechanisms that produce them (Cunningham et al., 2017; Moore & Benowitz, 2019). Following this logic, we hypothesized that the gene *takeout*, which has been shown to be involved in timing and feeding (Fujikawa et al., 2006; Guo et al., 2011; Meunier et al., 2007; Saito et al., 2006; Sarov-Blat et al., 2000; So et al., 2000; Vanaphan et al., 2012), may have been co-opted to enable the evolution of temporal kin recognition. We found that the version of *takeout* we examined was associated with the likelihood that both females and males would accept their larvae, such that beetles with higher *takeout* expression were more likely to accept their larvae. Somewhat contrary to our a-priori expectations, *takeout* was not associated with hunger or a transition to the full behavioural repertoire associated with parenting.

Co-option as an evolutionary process predicts that pre-existing mechanisms are transferred to a different context for a new function (Plachetzki & Oakley, 2007; West-Eberhard, 2003). For this reason, it was surprising that *takeout*, which has been shown to integrate timing with feeding (Sarov-Blat et al., 2000), was associated with temporal kin recognition but not hunger state in *N. orbicollis*. One explanation is that a different form of *takeout* influences feeding in *N. orbicollis*. While our data support placing the gene we examined in the *takeout* family, *takeout* typically has multiple copies (Vanaphan et al., 2012). Alternatively, it is possible that our methods were not sufficient to indicate an association between hunger and *takeout*. This seems unlikely, as similar procedures successfully induced starvation and reduced body mass in *Nicrophorus* and other comparably sized beetles (Auerswald & Gäde, 2000; Trumbo & Robinson, 2004; Trumbo & Xhihani, 2015) and removing food produced variation in *takeout* mRNA within ten hours in *Drosophila* (Sarov-Blat et al., 2000). We also did not find an association between *takeout* and development of parenting, i.e., days elapsed since egg laying, suggesting that *takeout* is not associated with parental behaviour generally but is more specific to the acceptance of larvae. These data suggest that the version of *takeout* we examined may play a specific role limited to temporal kin recognition, consistent with the specificity it has with timing of feeding.

Parents varied in both their likelihood of accepting larvae that showed up at the correct time and in *takeout* expression. Female and male parents often differ in how precise they are in discriminating their offspring from others across taxonomic groups, from vertebrates to invertebrates (Pinxten et al., 1991; Ringler et al., 2016), and it is an open question why the sexes differ in this regard. On a proximate level, females and males often use different cues to “start the clock” for temporal kin recognition, which can produce an asymmetry in information such that females have equal or more precise information on when to transition to parenting than males do (Davies et al., 1992; Elwood, 1994). For example, female mice experience physiological changes over pregnancy, while in males, mating and social interactions with the female influence the likelihood males will transition to a non-infanticidal state (Elwood & Kennedy, 1991; Elwood & Stolzenberg, 2020; Mei et al., 2023; Perrigo et al., 1992; Perrigo et al., 1990). Since males are more likely to have less reliable information about when offspring should arrive, this may lead to greater variation in acceptance. On a more evolutionary scale, it is a common pattern that females show more canalized care, whereas males are more environmentally sensitive in their care behaviour, including *Nicrophorus* (Benowitz & Moore, 2016; Smiseth et al., 2005).

We also found that *takeout* expression differed between the females and males in the analysis of parents that accept versus commit infanticide of their larvae. It is intriguing that *takeout* is associated with temporal kin recognition for both sexes, despite the likely scenario that females and males use different cues to initiate the transition to accepting offspring. *takeout* often has male-biased effects on social behaviours like courtship in *Drosophila*, with sex-specific expression in the fat body surrounding the brain in males but not females (Dauwalder et al., 2002; Lazareva et al., 2007; Vanaphan et al., 2012). Non-sex specific effects of *takeout* are generally associated with timing and feeding (Bauer et al., 2010; McDonald & Rosbash, 2001; Meunier et al., 2007; Saito et al., 2006; Smith et al., 2016; So et al., 2000), where *takeout* expression often occurs in chemosensory centers like the antennae (Dauwalder et al., 2002; Sarov-Blat et al., 2000). Guo (2011) found that *takeout* expression in antennae and hind leg of both sexes was involved in modulating attraction and repulsion to conspecifics through regulation of peripheral olfactory sensitivity. Since we found an association between *takeout* and temporal kin recognition in both sexes, it is an intriguing question for future research where and how *takeout* expression is associated with transitions in behaviour between females and males.

Ethologists have long predicted that complex behaviours can evolve through co-option of pre-existing behavioural precursors (Tallamy, 1984; West-Eberhard, 2003). We have shown an association between a gene that influences timing and feeding in other contexts with temporal kin recognition. Given that we did not see an association between hunger states and the general transition to a parenting state, this study raises many questions about the mechanisms by which *takeout* influences temporal kin recognition and its evolutionary history.

## Supplementary Information Appendix I

Additional information about the quantitative real-time PCR protocol

I. Additional information suggested in the Minimum Information for Publication of Quantitative Real-Time PCR Experiments (MIQE) guidelines not already provided in this paper.

A. Quantitative real-time PCR (qRT-PCR) primer sequences

*takeout-*

1. forward: GGGACATGGTCCAGCAAATA
2. reverse: GCATAATCGGTACCTCCCTTT

*gapdh*-

1. forward: CGACTACATGGTGTACCTGTTC
2. reverse: TGCCGTTGACGATGAGTTT

Primers were manufactured by Integrated DNA Technology (IDT, Coralville, IA, USA) and purified with IDT’s standard desalting technique.

A. qRT-PCR validation

Primer efficiency:

*takeout:* E=1.96, r^2^=0.9993

*gapdh:* E=1.95, r^2^=0.9996

